# Multielectrode Network Stimulation (ME-NETS) demonstrated by concurrent tDCS and fMRI

**DOI:** 10.1101/2023.06.13.544867

**Authors:** David A. Ross, Anant B. Shinde, Karl D Lerud, Gottfried Schlaug

**Affiliations:** Department of Neurology, Baystate Medical Center, and University of Massachusetts Chan Medical School – Baystate Campus, Springfield, MA 01107; University of Massachusetts – Amherst, Institute for Applied Life Sciences and Department of Biomedical Engineering, UMass Amherst, Amherst, MA 01003

**Author notes:** Corresponding authors Gottfried Schlaug, David Ross.

## Abstract

Non-invasive transcranial direct current stimulation (tDCS) can modulate activity of targeted brain regions. Whether tDCS can reliably and repeatedly modulate intrinsic connectivity of entire brain networks is unclear. We used concurrent tDCS-MRI to investigate the effect of high dose anodal tDCS on resting state connectivity within the Arcuate Fasciculus (AF) network, which spans the temporal, parietal, and frontal lobes and is connected via a structural backbone, the Arcuate Fasciculus (AF) white matter tract. Effects of high-dose tDCS (4mA) delivered via a single electrode placed over one of the AF nodes (single electrode stimulation, SE-S) was compared to the same dose split between multiple electrodes placed over AF-network nodes (multielectrode network stimulation, ME-NETS). While both SE-S and ME-NETS significantly modulated connectivity between AF network nodes (increasing connectivity during stimulation epochs), ME-NETS had a significantly larger and more reliable effect than SE-S. Moreover, comparison with a control network, the Inferior Longitudinal Fasciculus (ILF) network suggested that the effect of ME-NETS on connectivity was specific to the targeted AF-network. This finding was further supported by the results of a seed-to-voxel analysis wherein we found ME-NETS primarily modulated connectivity between AF-network nodes. Finally, an exploratory analysis looking at dynamic connectivity using sliding window correlation found strong and immediate modulation of connectivity during three stimulation epochs within the same imaging session.

## Introduction

Non-invasive transcranial direct current stimulation (tDCS) is of increasing interest both as a research tool to probe causal relationships between brain regions and behavior (Filmer et al., 2014; Hunter et al., 2015; Javadi & Cheng, 2013; Weber et al., 2014) and as a therapeutic tool for treating neurologic and psychiatric disorders (Fridriksson et al., 2019; Gunduz et al., 2021; Mondino et al., 2014; Palm, Ayache, et al., 2016).

Brain mapping techniques such as functional magnetic resonance imaging (fMRI) have found clear evidence that tDCS can modulate neural activity, both at the stimulation site and in remote brain regions (Amadi et al., 2014; Antonenko et al., 2017; Hunter et al., 2015; Zheng et al., 2011; Zheng et al., 2016). Yet, the effect of tDCS on functional connectivity varies considerably across studies, with studies reporting increased connectivity (Amadi et al., 2014; Krishnamurthy et al., 2015; Sotnikova et al., 2017), decreased connectivity (Antonenko et al., 2017; Dedoncker et al., 2019; Ficek et al., 2018), mixed effects (Mondino et al., 2016; Shahbabaie et al., 2018), and even no effect (Dissanayaka et al., 2017). These differences may be explained by factors such as the polarity and placement of electrodes (Li et al., 2019), the brain state during stimulation or training (Li et al., 2019; Sotnikova et al., 2017), and the dose/duration of stimulation (Shinde et al., 2021).

As a therapeutic tool, tDCS is often combined with a task designed to activate the target brain network (Ficek et al., 2018; Vines et al., 2011). Unfortunately, this can be a challenge in itself. For example, when language/speech-motor functions are severely impaired after a stroke affecting left-hemisphere one theorized path to language/speech-motor recovery is for connected regions within the unaffected right hemisphere to lead the recovery (Marchina et al., 2022; Pani et al., 2016; Schlaug, 2018; Turkeltaub et al., 2011; Xing et al., 2016) and/or using tasks that engage an auditory-motor feedforward and feedback network on the right hemisphere (Marchina et al., 2022; Norton et al., 2009; Pani et al., 2016; Schlaug, 2018; Turkeltaub et al., 2011; Xing et al., 2016).

At its core the AF-network has a white matter fiber bundle consisting of direct and indirect fibers that bidirectionally connect the posterior middle/superior temporal, inferior parietal, and posterior inferior frontal regions. While the AF-network is typically larger and more elaborate in the left hemisphere (Catani & Mesulam, 2008; Vernooij et al., 2007; Yeh, 2020), its right-hemisphere homotop (1) contains the rudimentary components allowing feedforward and feedback control of vocal-motor operations (Guenther & Vladusich, 2012; Tourville & Guenther, 2011; Tourville et al., 2008), and (2) exhibits structural plasticity as well as functional connectivity changes during recovery from stroke (Marchina et al., 2022; Pani et al., 2016; Schlaug et al., 2008; Vines et al., 2011; Xing et al., 2016).

Here, we investigate whether connectivity between nodes of the right hemisphere AF- network can be manipulated using a novel high-dose tDCS (4 mA) montage and concurrent tDCS-MRI in the absence of a behavioral task/training. Just as introducing a task can bias the effect of tDCS towards enhancing connectivity between brain regions involved in the task, we explored whether simultaneously applying tDCS to multiple distant but structurally connected nodes the AF-network might enhance connectivity by promoting a brain state favoring connectivity between those nodes. Specifically, we test (1) whether a multi-electrode montage simultaneously delivering stimulation to cortical nodes of the right AF-network (ME-NETS) enhances functional connectivity between those nodes more reliably than a single electrode delivering the same total current to just one node (SE-S), and (2) if changes in connectivity are specific to the targeted AF-network.

## Methods

### Subjects

A total of 23 subjects (age: *M* = 30.7 years, *SD* = 13.8 years; 13 female) were recruited through posted flyers and word of mouth from sites at Beth Israel Deaconess Medical Center (BIDMC, n = 9) and University of Massachusetts Amherst (UMass Amherst, n = 14). All subjects were right-handed according to the Edinburgh Handedness Inventory (Oldfield, 1971). Subjects were screened for a history of neurologic or psychiatric conditions, contra-indications to MRI and tDCS (e.g., implanted electric, metallic, or magnetic material) and additional contra-indications to tDCS (e.g., seizure disorder, scalp lesions where electrodes were supposed to be placed). Finally, most subjects were naïve to tDCS and underwent a familiarization session where they received a brief exposure to 4 mA tDCS in the mock scanner. A few subjects had participated in other studies and did not require that kind of a familiarization session. Subjects were randomly assigned to one of 3 groups which then determined their follow-up studies. No electrodes were mounted for the No-Stimulation session, which was otherwise of the same length as the concurrent tDCS-fMRI sessions. The study was approved by the institutional review boards at Beth Israel Deaconess Medical Center and the University of Massachusetts Amherst, and all subjects provided written informed consent.

At recruitment, subjects were asked if they could participate in up to 3 tDCS-MRI sessions, conducted on different days (though not necessarily consecutively). The intention was for subjects to participate in all three stimulation conditions (NS, SE, ME). However, due to the shutdown of non-clinical research studies during the COVID-19 pandemic, the lab relocating from BIDMC to UMass Amherst, and people leaving the area there were a number of dropouts. The final subject count in each condition was n=18, 15, and 13 for NS, SE, and ME respectively.

### Electrode Placement

Prior to entering the scanner room to participate in one of the tDCS-fMRI sessions, subjects were fitted with an electrode montage. For SE sessions, subjects were assigned one of three possible montages, depending on which AF-network node was to receive stimulation (n=5, 6, and 4 for the IFG, STG/MTG, and SMG nodes respectively). An anodal electrode (diameter = 4 cm) was then affixed to the scalp at the skull position of the assigned node. For ME sessions, there was one electrode montage comprising three anodal electrodes (diameter = 3 cm) affixed to the scalp at skull positions corresponding to each of the right-hemisphere AF-network nodes. In both conditions, a cathodal electrode (diameter 5 cm) was placed over the left supraorbital region.

Since our goal was to compare the effect of anodal stimulation over a single AF- network node with the effect of anodal stimulation over multiple AF-network nodes, we used the same total current of 4 mA for both the SE and ME condition. For the SE montages, the current was delivered via one round rubber electrode, giving a total (or max) charge density of 0.2292 C/cm2. For the ME montage, the 4mA current was split between three round rubber electrodes, with the STG/MTG electrode delivering 2 mA and the IFG and SMG electrodes each delivering 1 mA, giving a total charge density of 0.1019 C/cm2 across all three electrodes and a max charge density of 0.2132 C/cm2 at the STG/MTG electrode that got the highest current.

We used the 10-20 EEG system to determine electrode positions. The right IFG electrode was located at half the distance between C6 and F8, the right SMG electrode was located at one third the distance between C6 and CP6, and the right STG/MTG electrode was located half the distance between CP6 and TP8 (Figure 1). The left supraorbital cathodal electrode was located at FP1. Before placing an electrode on a subject’s scalp, the site was wiped clean with alcohol pads and as much hair was moved out of the way as possible to maximize contact and thus electrical conductance between electrode and skin. The electrodes (2 mm thickness) were then lathered with approximately 2 mm of Ten20 conductive paste and pressed onto the scalp. A self-adhesive bandage and hypoallergenic medical tape were used to hold the electrodes in place and electrode connectors were adjusted to avoid crossover between wires.

**Figure 1.**
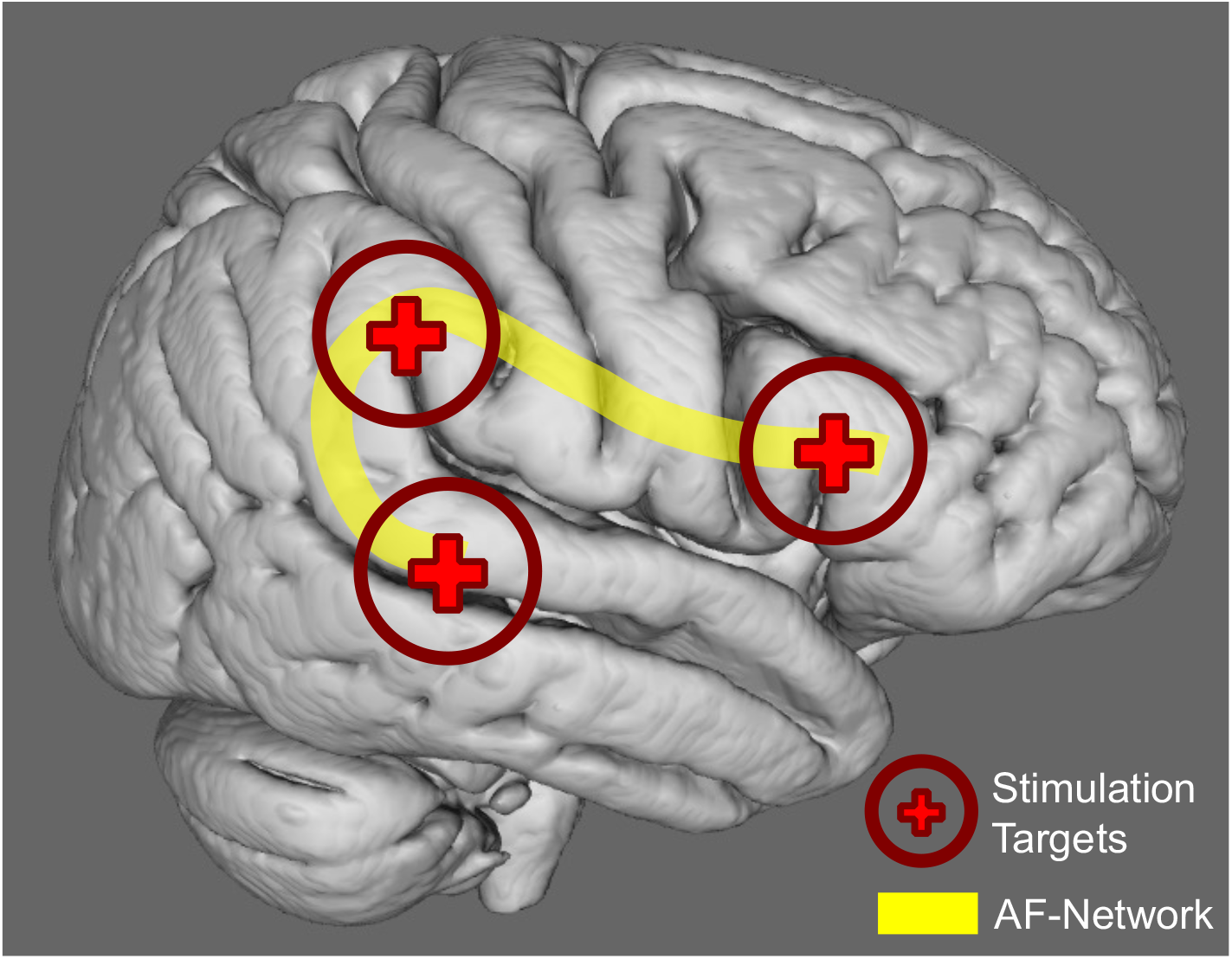
Schematic of Electrode Placements in Relation to the Arcuate Fasciculus (AF) The path of the right-hemisphere Arcuate Fasciculus (AF) is shown in yellow, passing through the parietal lobe and connecting posterior temporal gyrus to inferior frontal gyrus. The position of the three electrodes is also shown with sites over pTG, SMG, and IFG.

### fMRI-tDCS Procedure

Subjects were positioned in the scanner in a supine position. Foam padding was placed around their head to minimize movement and earplugs were provided to minimize any discomfort caused by MRI noise. For tDCS-fMRI sessions (SE, ME), the scalp electrodes were connected to an MR compatible direct-current multi-channel (DCMC) stimulator located outside of the scanner room (NeuroConn, NeuroCare Group, Germany).

During every session, we collected a low resolution T1w scan and a 24-minute resting-state scan. The low resolution T1w scan was collected at the start of the session and was inspected to confirm that the electrodes were correctly positioned over the intended anatomical region for tDCS-fMRI scans. If the electrodes needed to be moved, the subject was taken out of the scanner, the electrodes were repositioned, and the low resolution T1w scan was repeated. High-resolution T1w and T2w scans were also collected for all subjects, typically at the end of the first scanning session, and were used to model the electric field.

For tDCS-fMRI sessions, scalp electrodes were removed before collecting the high-resolution scans to get the best estimate of scalp thickness. Finally, for some subjects we collected Diffusion Tensor Imaging (DTI) data (before and after the resting-state scan) and a second resting-state scan (immediately following the first resting-state scan). These data were not analyzed for the current study and will be published separately.

During the functional imaging scans, subjects were instructed to keep their eyes open and fixate on a small green crosshair that was displayed on a screen visible via the head coil-mounted mirror. For tDCS-fMRI scans (SE, ME), stimulation was applied during 3 equally spaced 4-minute epochs starting at 3, 10, and 17 minutes. The stimulation signal was generated by the MR compatible DCMC stimulator, converted to a box cable compatible signal, and relayed via a CAT5-DB9 adapter cable into the scanner room. On the other side of the RF panel, a cable relayed the signal to an adapter box where it was converted into a stimulation signal and delivered via the rubber scalp electrodes. To reduce the chances of subjects experiencing an uncomfortable sensation at the electrode sites, the current was gradually ramped up over the first 30 seconds of each tDCS epoch from 0 mA to 4 mA. The current was also gradually ramped down to 0 mA over the last 30 seconds of each tDCS epoch, to avoid sudden transitions that might increase subject movement or discomforts.

After tDCS-fMRI sessions, we checked the subject’s scalp for skin burns or other lesions and asked them to indicate their experience of the tDCS stimulation from 0-10 on a visual analog scale (0 indicated that subjects tolerated the stimulation well and had no unusual sensations and a score of 10 was an intolerable experience). None of the subjects reported scores higher than 8.

### MRI Parameters

Subjects were scanned on either a 3-Tesla (3T) GE Discovery MR750 scanner located at BIDMC or a 3T Siemens Skyra scanner located at UMass Amherst, depending on where they were recruited. All images were acquired with 32-channel head coil. Note that scan sequences were similar at the two sites and a preliminary analysis of the data (reported in the Results section) did not show systematic differences between study sites.

#### BIDMC MR Scan Details

T1w anatomical images were acquired using a 3D fast SPGR (BRAVO) sequence from GE in the sagittal plane (Low-resolution T1w scan parameters: TR = 7.0 ms, TE = 2.6 ms, flip angle = 12°, FOV = 240 mm, matrix size = 256 x 256, 96 slices, voxel size = 2.0 x 0.94 x 0.94 mm^3^; High-resolution T1w scan parameters: TR = 8.2 ms, TE = 3.2 ms, flip angle = 12°, FOV = 240 mm, matrix size = 256 x 256, 180 slices, voxel size = 1.00 x 0.94 x 0.94 mm^3^). Resting state functional scans were acquired using gradient-echo echoplanar imaging in the axial plane (TR = 3196 ms, TE = 24 ms, flip angle = 90°, FOV = 240 mm, matrix size = 128 x 128, 58 slices, and voxel size = 1.88 x 1.88 x 2.5 mm^3^). High-resolution T2w images were acquired using a 3D FSE CUBE sequence from GE in the sagittal plane (TR = 5002 ms, TE = 98.6 ms, flip angle = 90°, FOV = 240 mm, matrix size = 512 x 512, 180 slices, and voxel size = 1 x 1 x 1 mm^3^).

#### UMass Amherst MR Scan Details

T1w anatomical images were acquired using a 3D IR sequence in the sagittal plane (Low-resolution T1w scan parameters: TR = 1490 ms, TE = 3.4 ms, flip angle = 9°, FOV = 256 mm, matrix size = 256 x 256, 96 slices, and voxel size = 2.0 x 1.0 x 1.0 mm^3^; High-resolution T1w scan parameters: TR = 2020 ms, TE = 3.4 ms, flip angle = 9°, FOV = 256 mm, matrix size = 256 x 256, 208 slices, and voxel size = 1.0 x 1.0 x 1.0 mm^3^). Resting state functional scans were acquired using gradient-echo echoplanar imaging in the axial plane (TR = 3000 ms, TE = 31 ms, flip angle = 90°, FOV = 210 mm, matrix size = 84 x 84, 60 slices, and voxel size = 2.5 mm^3^). High-resolution T2w images were acquired using a 3D SE sequence in the sagittal plane (TR = 3200 ms, TE = 408 ms, flip angle = 120°, FOV = 256 mm, matrix size = 256 x 256, 208 slices, and voxel size = 1 mm^3^).

### Preprocessing and Denoising

MRI data were converted to NIfTI format using dcm2niix (Li et al., 2016) before undergoing preprocessing and denoising in the CONN toolbox (Whitfield-Gabrieli & Nieto-Castanon, 2012): Functional images were realigned using the SPM12 realign and unwarp procedure (Andersson et al., 2001); outlier scans were identified using CONN’s implementation of ART toolbox with the default settings (framewise displacement > 0.9 mm and/or global BOLD signal change > 5 SDs; (Mazaika et al., 2009); functional and anatomical images were normalized to MNI space and segmented into GM, WM, and CSF tissue classes using the SPM12 unified segmentation and normalization procedure (Ashburner & Friston, 2005); and finally the functional images were smoothed by spatial convolution with a Gaussian kernel of 8mm full width half maximum.

Functional denoising was carried out in the CONN toolbox using Ordinary Least Squares regression. Potential confounding factors included motion parameters and their 1st order derivatives (12 factors), outlier scans (a maximum of 66 factors), white matter (5 factors) and CSF (5 factors) noise components estimated using the CONN’s implementation of aCompCor (Behzadi et al., 2007), and a linear detrending term. In addition, stimulation condition effects (On, Off) and their 1st order derivatives (2 factors) were included as confounding factors to remove any constant tDCS induced responses (Nieto-Castanon, 2020). Specifically, we wanted to identify the modulatory effect of tDCS on connectivity and not simply the direct effect of tDCS on the BOLD signal (e.g., in two simultaneously stimulated brain regions). Finally, further controlling for the direct effect of tDCS on BOLD signal and to control for other high- and low-frequency confounds the functional timeseries were band- pass filtered to remove frequencies below 0.008 Hz or above 0.09 Hz using CONN’s ‘Simult’ option to implement simultaneous filtering and regression (Hallquist et al., 2013; Nieto-Castanon, 2020).

### Region of Interest (ROI) Definitions

To explore connectivity within the right-hemisphere AF-network, we created sphere ROIs (8 mm radius) at the cortical locations we had targeted with anodal stimulation. Two ROIs were defined for each AF-network node (Figure 2A): SMG (aSMG, pSMG), posterior temporal gyrus (pMTG, pSTG), and IFG (IFG tri, IFG oper). The central coordinates for each ROI were taken from CONN’s default atlas (center of gravity determined using SPM12), but we chose to use spheres to avoid possible confounds related to differences in ROI size. In addition, to explore how specific any effect of tDCS was to the targeted network we created a second set of sphere ROIs (8 mm radius) located at cortical regions connected by the right- hemisphere Inferior Longitudinal Fasciculus (ILF; (Latini et al., 2017; Panesar et al., 2018). As we did for the AF-network, we created 6 ILF-network ROIs using coordinates taken from CONN’s default atlas (Figure 2B). Three ROIs were located in posterior temporal and occipital lobe: inferior lateral occipital cortex (iLOC), temporal occipital fusiform gyrus (TOFusG), and lingual gyrus (LG). And three ROIs were located in anterior temporal lobe: anterior inferior temporal gyrus (aITG), anterior temporal fusiform gyrus (aTFusG), and temporal pole (TP).

**Figure 2.**
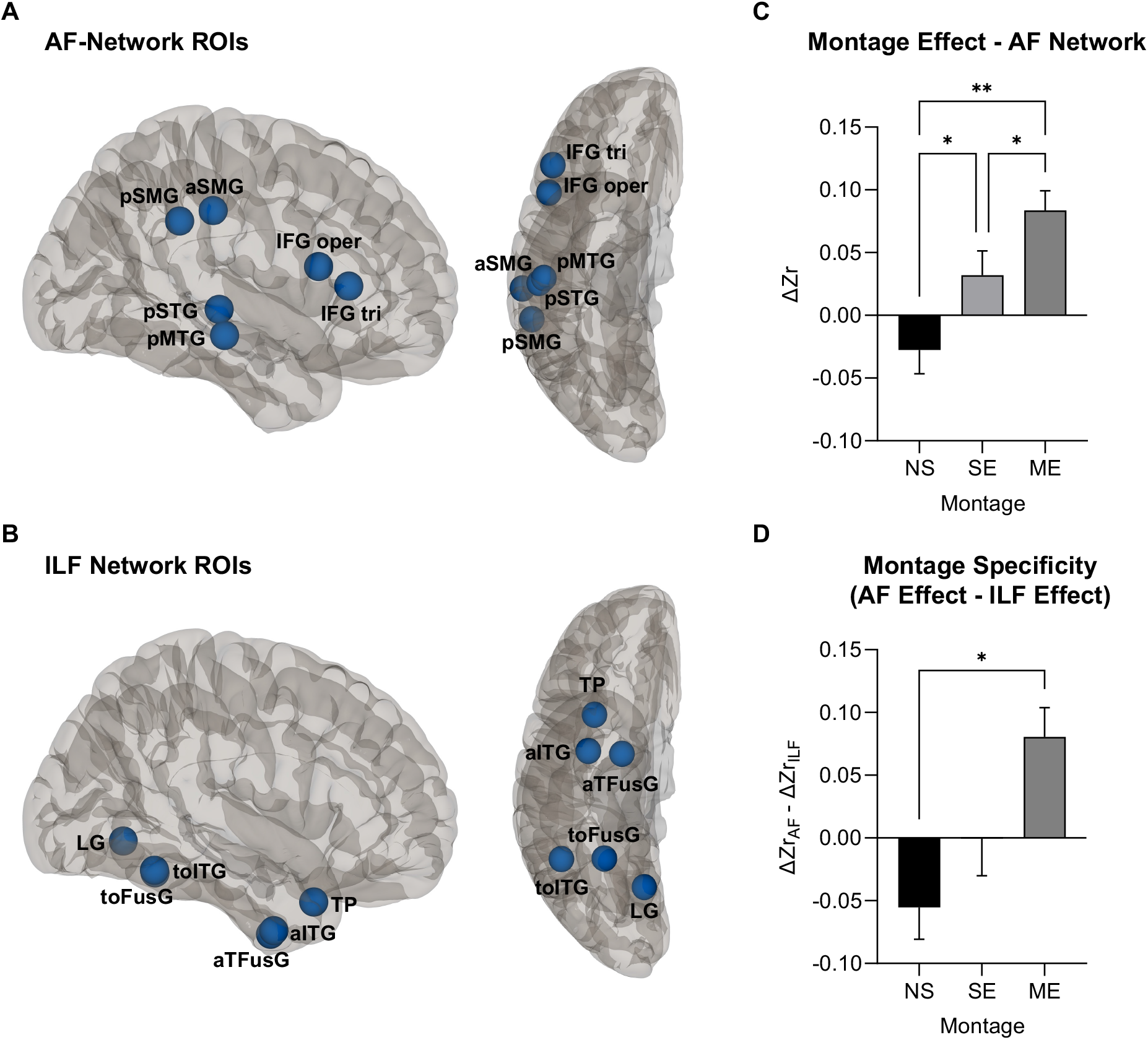
ROI-to-ROI Analysis: ROIs Placement and Montage Effect for the AF- and ILF- Networks. ROI placement and results for the ROI-to-ROI analyses. ROI placement is shown on the left: (**A**) the position of right-hemisphere AF-network ROIs; (**B**) the position of right-hemisphere ILF-network ROIs. The effect of montage – no stimulation (NS), single electrode (SE), and multi electrode (ME) – on connectivity is shown on the right: (**C**) The effect of tDCS montage on connectivity within the AF-network; (**D**) The specificity of the tDCS effect, defined as the difference between the effect of a montage on AF-network connectivity and its effect on ILF-network connectivity. Asterisks indicate significant differences: p < .05 (*) and p < .01 (**).

### ROI-to-ROI Analyses

To investigate the effect of montage (NS, SE, ME) on AF-network connectivity we extracted the connectivity (Fisher transformed correlations, Zr) between seed ROIs in the SMG (aSMG, pSMG) and posterior temporal gyrus (pMTG, pSTG), and target ROIs in the IFG (IFG tri, IFG oper). For each subject and montage (NS, SE, ME) we calculated the average difference in connectivity across AF-network ROIs between stimulation ON and stimulation OFF (ΔZr). Similarly, we computed the effect of montage on ILF-network connectivity, calculating the difference in connectivity between stimulation ON and stimulation OFF (ΔZr) between seed ROIs in posterior temporal and occipital lobe (iLOC, TOFusG, LG) and target ROIs located in anterior temporal lobe (aITG, aTFusG, TP). Note, the NS condition was analyzed in the same way as SE and ME, treating the 4-minute epochs starting at 3, 10, and 17 minutes as a pretend stimulation ON and the rest of the timecourse as a pretend stimulation OFF.

To analyze ROI-to-ROI data, we implemented a mixed model using GraphPad Prism 9.0. This allowed us to correctly account for our repeated measures design despite missing data (i.e., not all subjects ended up completing all conditions). GraphPad Prism’s implementation of a mixed model uses a compound symmetry covariance matrix and is fit using Restricted Maximum Likelihood (REML). The results of this implementation can be interpreted like the results of a one-way repeated measures ANOVA, and in the absence of missing data it would give identical P values. Degrees of freedom were adjusted using the Geisser and Greenhouse epsilon hat method to account for potential violations of sphericity.

### Seed-to-Voxel Analysis

To further explore the spatial extent of tDCS modulated connectivity, we computed connectivity between the right AF-network nodes (IFG tri, IFG oper, pSTG, pMTG, aSMG, pSMG) and the rest of the brain. Using CONN’s General Linear Model framework we tested for an effect of tDCS (On > Off) on connectivity between each AF ROI and the rest of the brain. Separate contrasts were computed for each montage (SE, ME) and ROI (equivalent to paired t-tests). We used CONN’s default thresholds for parametric statistics based on Random Field Theory (Worsley et al., 1998): voxel threshold p<0.001, and family-wise error (FWE) corrected cluster threshold (p-FWE<0.05), with Bonferroni correction applied to the FWE threshold to account for multiple comparisons for each seed, p-FWE / 12 (6 seeds x 2 montages).

### Sliding-Window Correlation Analysis

For each subject and montage (NS, SE, ME), preprocessed time series were extracted from the four AF-network seeds in SMG (aSMG, pSMG) and posterior temporal region (pMTG, pSTG), and the two target ROIs in the IFG (IFG tri, IFG oper). To account for the slight difference in TR between scans acquired at UMass Amherst (TR=3000 ms) and those acquired at BIDMC (TR=3169 ms), time series were resampled to a resolution of 3000 ms using the linear interpolation method in MATLAB. Sliding-window correlation time series were then computed for each seed-target pair by calculating the Fisher transformed correlation (Zr) between the two time-series using a 30 TR (90 s) window and a step size of 1 TR.

## Results

### Montage Effect within the AF-Network

Our primary hypothesis was that high-dose tDCS delivered to multiple nodes of the AF-network (ME) would modulate connectivity between nodes more strongly than the same total dose delivered to a single node (SE). In addition, we also compared each of the stimulation conditions (SE, ME) to a no-stimulation baseline (NS). This was important because subjects all received tDCS on the same schedule (i.e., the same stimulation onset times), thus we compared our effects to a NS baseline to rule out the possibility that our effects might simply be due to systematic effects of time-in-scanner on connectivity. To ensure that our results were not biased by Study Site, we first ran a two-way mixed model (equivalent to a two-way ANOVA) with factors Study Site (UMass Amherst, BIDMC) and Montage (NS, SE, ME). This analysis revealed a main effect Montage (*p* = .0052) but no main effect of Study Site (*p* = .4793) or Study Site x Montage interaction (*p* = .4930). Therefore, we collapsed across Study Site for all subsequent analyses. In line with our hypotheses, a one-way mixed model revealed a significant effect of montage (NS, SE, ME) on connectivity (ΔZr) within the AF-network, *F*(1.89, 40.60) = 9.16, *p* = .001, ε = .94 (Figure 2C). Moreover, planned comparisons revealed that ME stimulation increased AF-network functional connectivity more than SE stimulation, *t*(9)=2.39, *p* = .0404, with both ME and SE stimulation increasing functional connectivity significantly compared to the NS baseline, ME vs. NS: *t*(9) = 3.83, *p* = .004; SE vs. NS: *t*(11) = 2.50, *p* = .0298.

### Specificity of the Montage Effect

To investigate whether the effect of tDCS on connectivity was specific to the targeted network, we compared the effect of each montage on connectivity within AF-network (Figure 2C) to the effect of each montage on connectivity within the ILF-network. Within ILF- network there was no effect of Montage on connectivity (ON-OFF), *F*(1.96, 42.13) = 0.47, *p* = .63, ε = .98, and one-sample t-tests indicated that there was no significant effect of stimulation on connectivity for any Montage: NS (*M* = 0.03), *t*(17) = 1.49, *p* = 0.1533; SE (*M* = 0.02), *t*(14) = 1.28, *p* = 0.2213; ME (*M* = 0.00), *t*(12) = 0.12, *p* = 0.9054. To test for an interaction effect between Montage (NS, SE, ME) and Network (AF, ILF) we entered the difference score (ΔZrAF - ΔZrILF) into a mixed model revealing a significant effect of montage (NS, SE, ME) on AF-network specific connectivity (Figure 2D), *F*(1.99, 42.70) = 6.43, *p* = .0037, ε=.99. Planned comparisons revealed that ME stimulation increased AF-network connectivity more than ILF-network connectivity compared to the NS baseline, *t*(9) = 3.18, *p*=.0111, but SE stimulation did not, *t*(11) = 1.71, *p* = .1152.

### Seed-to-Voxel Results

We conducted a seed-to-voxel analysis to investigate the effect of tDCS (On - Off) on connectivity between nodes of right hemisphere AF-network (IFG tri, IFG oper, pSTG, pMTG, aSMG, pSMG) and the rest of the brain. For ME but not SE, we found evidence for several regions where connectivity with an AF-network node was significantly modulated by stimulation (Figure 3; Table 1). In line with the findings from the ROI-to-ROI analysis, the majority of clusters were around AF-network nodes, namely right posterior temporal gyrus (pMTG, pSTG), right supramarginal gyrus (aSMG, pSMG), and right inferior frontal gyrus (IFG tri, IFG oper). Remarkably, very few clusters lay outside of the AF-network (paracingulate gyrus, right lingual gyrus, Cuneal, and cerebellum) and most clusters were in the right-hemisphere (with the exception of the paracingulate and cerebellar cluster), suggesting that ME stimulation was able to modulate connectivity in a very targeted way. For the SE condition we found a single cluster in right frontal pole where stimulation modulated connectivity with IFG.

**Figure 3.**
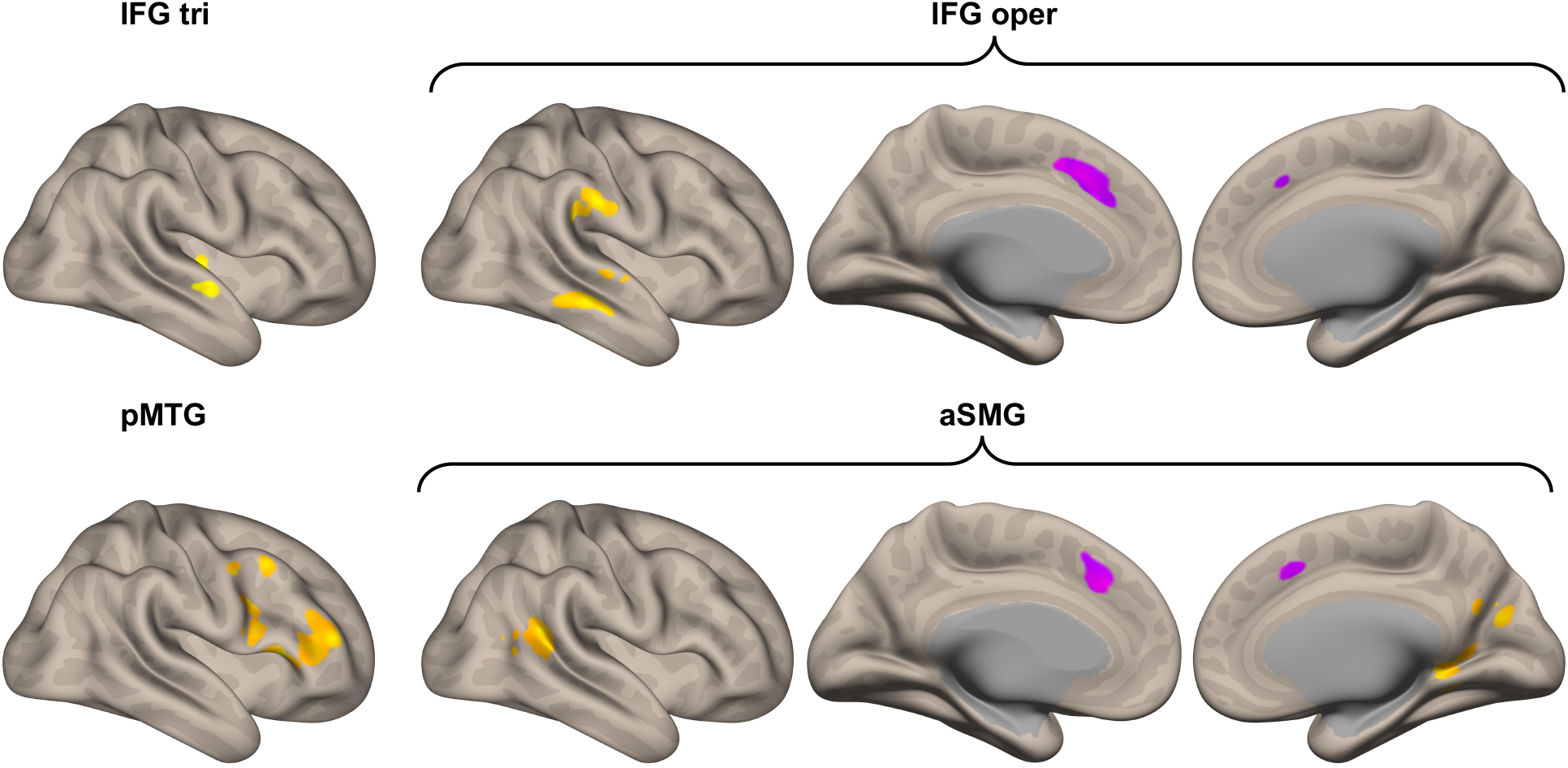
The Effect of ME tDCS on Connectivity Between the AF-Network and the Rest of the Brain. Highlighted brain regions indicate where ME tDCS significantly modulated connectivity (ON – OFF) with one of the AF-network seeds (IFG tri, IFG oper, pSTG, pMTG, aSMG, pSMG). Regions where stimulation increased connectivity with AF-network (ON > OFF) are shown in orange – primarily right-hemisphere posterior temporal, parietal, and inferior frontal regions connected by AF white matter tract. Regions where stimulation decreased connectivity with AF-network (ON < OFF) are shown in purple – primarily the left- hemisphere paracingulate region. Clusters were defined with voxel threshold p < .001 and cluster threshold p-FWE < .05, which was Bonferroni corrected for multiple comparisons p- FWE / 12 (6 seeds x 2 montages). Seeds are not shown here if there were no significant clusters (pSTG, pSMG).

**Table 1.**
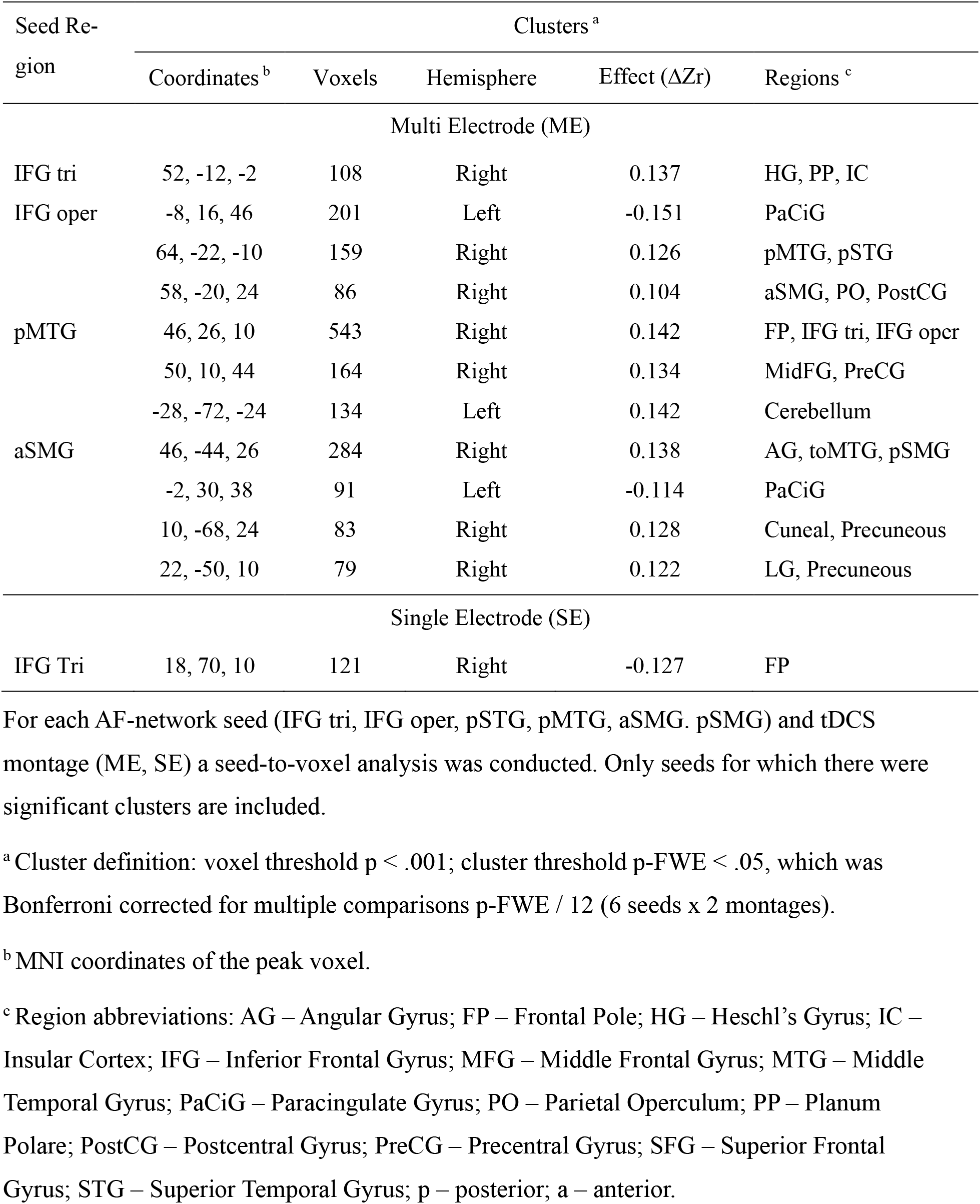
Clusters where tDCS Significantly Modulated Connectivity with the AF-network

### Sliding-Window Correlation Results

In addition to our primary static-connectivity analyses, we conducted an exploratory sliding- window correlation analysis to capture dynamic connectivity and test and show in a graphic manner whether AF-network connectivity can be modulated reliably and repeatedly in a differential way by the two tDCS montages (SE, ME). The sliding-window correlation analysis provides a means of visualizing fluctuations in connectivity without making strict assumptions about their latency and duration relative to each bout of tDCS. The results of the sliding window correlation analyses are shown in Figure 4. For the ME condition (Figure 4A), peaks are clearly visible in the connectivity time course during each of the three ON epochs with corresponding troughs during the subsequent OFF epochs. In contrast, there do not seem to be any well-defined peaks in the SE (Figure 4B) or NS (Figure 4C) conditions.

**Figure 4.**
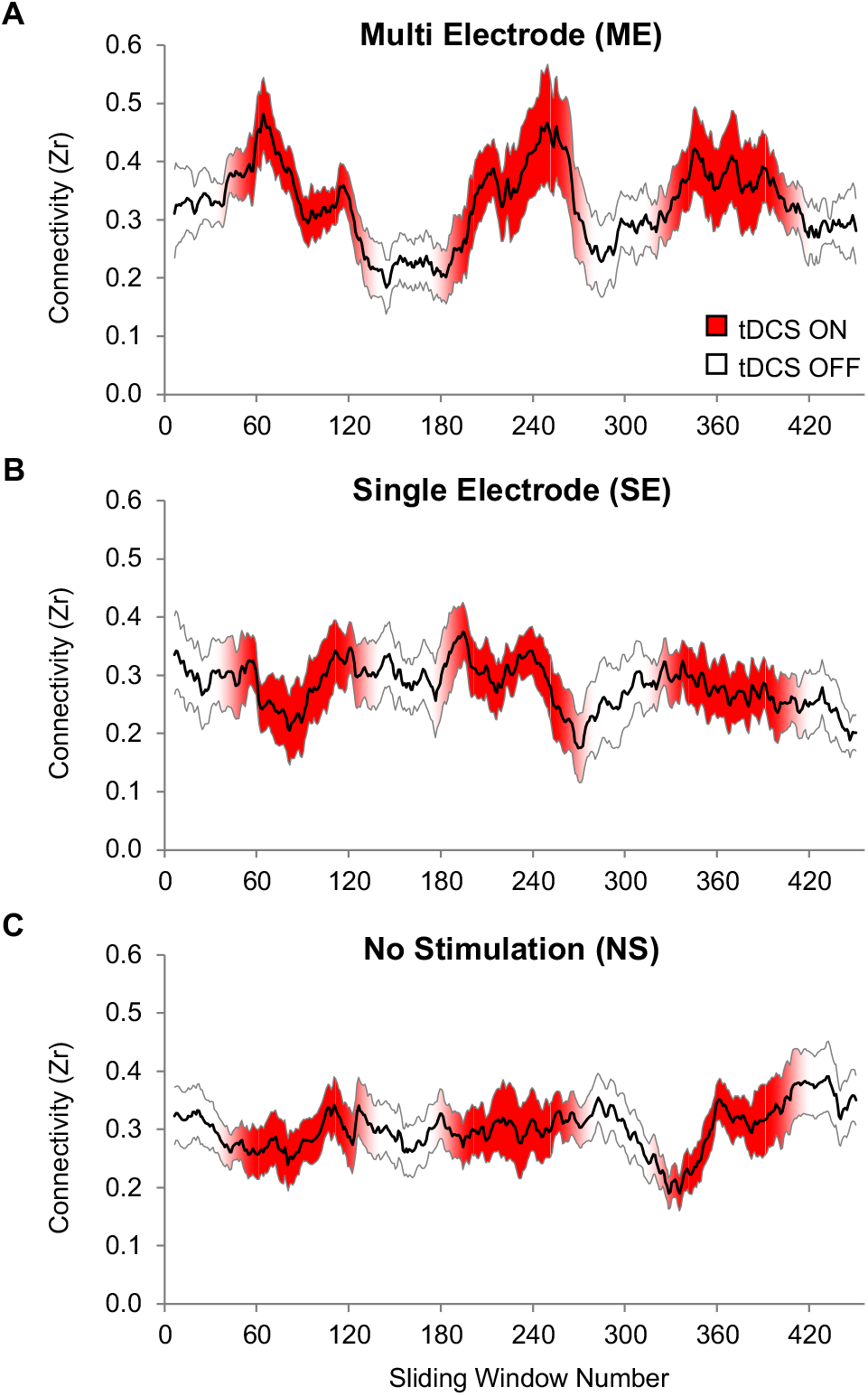
Dynamic Connectivity Within the AF-Network During tDCS-fMRI Sessions. Sliding window correlations (SWC) between nodes of the AF-network are shown for each montage: (**A**) ME, (**B**) SE, and (**C**) NS. Because each sliding window includes multiple timepoints (30 TRs, 90 sec) the onset and offset of each tDCS epochs is blurred: Solid red indicates segments of the SWC timeseries where the sliding window captured only timepoints from tDCS ON epochs; Solid white indicates segments of the SWC timeseries where the sliding window captured only timepoints from tDCS OFF epochs; Transition periods, where the sliding window includes both tDCS ON and tDCS OFF timepoints, are indicated by the linear transition between white and red. The solid black line indicated the mean with the grey lines showing the SE of the mean at each timepoint.

## Discussion

We found that multi electrode network stimulation (ME-NETS) differentially increased functional connectivity between nodes of a target network (the AF network) relative to a no stimulation (NS) baseline. Moreover, ME-NETS increased connectivity withing the target network more than single electrode stimulation (SE-S) with the same current (4 mA). It is quite novel to demonstrate such control over connectivity within a targeted network in the absence of task or without engaging the network in longer-term training exercises.

Results of both the ROI-to-ROI analysis and the seed-to-voxel analysis suggest that the changes in connectivity brought on by ME-NETS were relatively localized to the targeted AF network. In the case of the ROI-to-ROI analysis we found that ME-NETS modulated connectivity between AF-network nodes significantly more strongly than between nodes of the right-hemisphere ILF-network. Further supporting the idea that effects were spatially limited to the AF-network, most of the clusters that we uncovered in the seed-to-voxel analysis were within the AF-network.

The fact that changes in connectivity were largely isolated to the target network is particularly surprising given that these regions are often thought of as hubs with multiple diffuse patterns of functional connectivity depending on cognitive tasks (Fedorenko et al., 2013; van den Heuvel & Sporns, 2013). It might be expected that delivering tDCS stimulation to these nodes would simply enhance underlying patterns of connectivity, increasing connectivity between the stimulated region and all other regions to which it is connected. Or, since we delivered tDCS in the context of a resting state scan we might expect to see increases in functional connectivity between the stimulated region and regions associated with resting state such as the default mode network (Amadi et al., 2014; Grami et al., 2022; Keeser et al., 2011; Li et al., 2019). Indeed, while increased functional connectivity within spatially localized, task-specific networks has previously been reported, these effects have only been found in the presence of a task that targets that network (Callan et al., 2016; Hampstead et al., 2014; Krishnamurthy et al., 2015; Li et al., 2019).

Single-anodal-electrode modulation of functional connectivity between stimulated regions has been explored on short time scales, e.g. within a session during concurrent settings, as well as comparing before and after stimulation in interleaved settings (Amadi et al., 2014; Antonenko et al., 2017; Dedoncker et al., 2019; Ficek et al., 2018; Krishnamurthy et al., 2015; Meinzer et al., 2012; Mondino et al., 2016; Neeb et al., 2019; Sehm et al., 2012; Shahbabaie et al., 2018; Sotnikova et al., 2017) as well as on time scales of weeks over multiple sessions (Dunlop et al., 2015; Lefebvre et al., 2017; Palm, Keeser, et al., 2016). Results have been mixed, were typically of small magnitude, or had unexpected effects in either increasing or decreasing functional connectivities.

There are other multi-electrode techniques, such as HD-tDCS (Borckardt et al., 2012; Lefebvre et al., 2019; Nikolin et al., 2019; Richardson et al., 2014; Villamar et al., 2013; Yaqub et al., 2018; Zito et al., 2015), which differs from the ME-NETS technique in several ways. For one, there are usually multiple cathodal electrodes with only a single anode, though there are some studies that have experimented with multiple anodes (Richardson et al., 2014; Zito et al., 2015). Related to this, the main difference is that HD-tDCS is usually intended to increase focality and biological effects to a single location, whereas ME-NETS is intended to modulate activity in a wide network by distributing electrodes across a network of nodal target regions.

Our experimental approach also distinguishes the present study from other concurrent or interleaved tDCS-MR imaging studies. One of our aims was to show that functional connectivity can be modulated reliably and repeatedly, basically at the will of the experimenter. As a result of this experimental approach, one can see that functional connectivity modulation is immediate upon onset of the first bout of tDCS, and reliably increases during stimulation, and decreases during periods of no stimulation. Increased functional connectivity among this cluster might be related to the tDCS increasing blood flow and metabolic activity in the nodal cortical regions which then may lead to more coherence between these regions. Specifically, this increased coherence between these nodal regions could be triggered by current mainly flowing through the structural backbone of the network, the AF, and thus lead to more electrical and chemical communication between these regions.

Several methodological and translational questions arise from these results. This increased functional connectivity within targeted nodal regions within one hemisphere and with homotop regions on the other hemispheres may be a pathway to increasing autonomous communication between these areas over time, in particular if the stimulation might be paired with behavioral interventions that target the right AF network and as such could aid recovery from pathology as well as boosting brain function of the targeted network and system. A couple of experimental parameters could be further tested and explored in the future in ways that were not tenable in the present work. For one, the dosing at the three ME electrodes was 2-1-1 mA, corresponding to STG/MTG, SMG, and IFG respectively. The combined 4 mA dose has been found to be safe considering previous literature (Chhatbar et al., 2016) as well as our own experiences now, but it should be explored whether a different current distribution between the three electrodes could lead to even stronger effects than those observed in this study.

From a translational perspective, the main opportunity for the future is exploring behavioral correlates of this functional connectivity increase and to determine if adding a behavioral task concurrently might enhance the effects even more. The AF network on either hemisphere is largely involved in feedforward and feedback control of vocal-motor output, constituting an example of a redundant bihemispheric network that can subserve vocal communication purposes (Schlaug, 2018). The clinical application that seems most intuitive is recovery from large left hemisphere lesions caused by stroke or other brain injury. In many cases, such patients need to rely on the functioning and plasticity of the homotop networks on the right side (Marchina et al., 2022; Pani et al., 2016; Wan et al., 2014) and therapies based on ME-NETS in the right hemisphere may lead to further improvement and recovery of what is happening naturally already.

